# Cellular and molecular basis of *Pf*Coronin function in artemisinin resistance in *Plasmodium falciparum*

**DOI:** 10.1101/2025.09.06.674515

**Authors:** Imran Ullah, Madeline A. Farringer, Malhar Khushu, Anna Y. Burkhard, Erica Hathaway, Bailey C. Willett, Sara H. Shin, Aabha I. Sharma, Morgan C. Martin, Oliver Harrigan, Kairon L. Shao, Jeffrey D. Dvorin, Daniel L. Hartl, Sarah K. Volkman, Selina Bopp, Sabrina Absalon, Dyann F. Wirth

## Abstract

Artemisinin-based therapies are central to malaria treatment, but their efficacy is threatened by the emergence of resistant *Plasmodium falciparum* strains. While the role of *Pfkelch13* mutations in clinical resistance is well established, recent reports from Africa implicate *Pfcoronin* mutations in treatment failure, adding to the complexity of resistance mechanisms. Here, we show that *Pf*Coronin, a non-essential actin regulator active during the ring-stage, facilitates efficient hemoglobin uptake. We demonstrate that resistance-associated *Pfcoronin* mutations disrupt *Pf*Coronin’s interaction with *Pf*Actin and its ring-stage localization, leading to impaired endocytosis and reduced hemoglobin acquisition. *Pf*Coronin and *Pf*Kelch13 function in distinct cellular regions; mutations in both converge to limit heme availability and artemisinin activation. Although *Pf*Coronin is dispensable for parasite viability, our findings demonstrate that *Pfcoronin* mutations reduce endocytosis and modulate artemisinin susceptibility during the clinically relevant ring stage—highlighting how non-essential, temporally restricted proteins can shape antimalarial drug response and resistance.

## Introduction

Rapidly acting and potent antimalarial drugs, such as artemisinin (ART) and others in its class, are critical components of current front-line treatments, and these drugs have contributed to yearly reductions in malaria mortality for the past two decades^1^. However, partial resistance to ART and artemisinin combination therapies (ACTs) threatens their efficacy^2,3^. ART treatment failures initially emerged in Southeast Asia, and were associated with mutations in the *Plasmodium falciparum kelch13* locus^4^. The mechanism of *Pfkelch13*-mediated ART resistance is well studied, and prior work has demonstrated reduced hemoglobin uptake in ART-resistant *Pfkelch13* mutant parasites^5,6^. ART is a sesquiterpene lactone with an endoperoxide bridge that must be cleaved to potentiate antimalarial activity. Typically, this activation occurs via heme^7^, likely derived via endocytosis and digestion of hemoglobin. Thus, the observed reduction in endocytosis in *Pfkelch13* mutants may decrease the availability of free heme, slowing the activation of ART and leading to ART resistance ^5,6^.

While resistance research has largely focused on essential genes such as *Pfkelch13*, emerging evidence suggests that alternative, underrecognized molecular determinants may also contribute to resistance phenotypes^8–11^. As part of our previous efforts to understand potential ART-resistance mechanism(s) in Africa, we demonstrated *Pfcoronin* to be a major driver of *in vitro* partial ART resistance in Senegalese isolates of *P. falciparum*^12,13^. Coronin proteins are widely conserved, and while many organisms have multiple *coronin* genes, *P. falciparum* and other unicellular pathogens encode just one^14,15^. Like other Coronin orthologs, *Pf*Coronin includes a WD40 domain. These domains are thought to facilitate protein-protein interactions and are involved in a wide range of cellular functions^16,17^. Coronin proteins have been associated with actin-binding activities in multiple organisms^18^; in the apicomplexan parasite, *Toxoplasma gondii*, *Tg*Coronin regulates the rate and extent of actin polymerization, and is required for endocytosis and membrane recycling^19^. In *P. berghei*, *Pb*Coronin is implicated in the regulation of gliding motility in sporozoites^20^. In *P. falciparum*, *in vitro* biochemical studies have confirmed that the WD40 domain of *Pf*Coronin binds actin and organizes F-actin filaments into parallel bundles^21^. In late asexual blood-stage parasites, *Pf*Coronin localizes to the parasite periphery, potentially linking actin filaments to the plasma membrane^21^; localization in earlier stages has not been previously assessed.

Despite being considered non-essential for overall parasite viability, we hypothesized that *Pfcoronin* may nevertheless play a stage-specific and previously underappreciated role in drug response. Here, we systematically characterize *Pf*Coronin’s molecular interactions in both ring-stage and late-stage parasites, and assess the impact of resistance-associated mutations on its abundance, subcellular localization, and involvement in hemoglobin uptake. In doing so, we seek to clarify how a non-essential, temporally restricted cytoskeletal regulator could serve as a critical mediator of ART resistance, and in turn, expand current perspectives on the molecular basis of parasite adaptation and therapeutic failure.

## Results

### Reduced interaction of *Pf*Coronin mutants with *Pf*Actin in rings

We previously showed that mutations in *Pfcoronin* confer *in vitro* ART resistance in Senegalese *P. falciparum* isolates, as measured by the ring-stage survival assay (RSA)^12^. Either a single G50E mutation (Senegalese isolate from Thiès) or paired R100K/E107V mutations (Senegalese isolate from Pikine) are sufficient for resistance^12,13^. *Pf*Coronin features a seven-bladed WD40 domain, with resistance-associated Gly50, Arg100, and Glu107 residues located in the first two blades. Throughout this study, we used the Pikine line as a model to dissect the molecular basis of *Pfcoronin*-mediated resistance. For interaction studies, we generated parasite lines with endogenous C-terminal spaghetti monster-hemagglutinin (smHA)-tagged^22^ *Pf*Coronin in both wild-type (WT) and R100K/E107V (mutant) backgrounds (Figure S1).

To identify *Pf*Coronin interaction partners, we performed anti-HA immunoprecipitation from both early ring (3–9 hours post invasion, hpi) and trophozoite (32–40 hpi) stages in smHA-tagged lines followed by LC-MS/MS (Figure S2A). Proteins detected in untagged control lines were excluded as background, and remaining candidates were normalized to *Pf*Coronin bait levels for comparison. High-confidence interactors were prioritized if consistently detected across all replicates at ≥5% of *Pf*Coronin abundance. (Tables S1–S2). For analysis of differential association, we considered proteins with >5-fold differences in abundance or *p* < 0.05 between WT and mutant (Figure S2B/Table S3 [3–9 hpi]; Figure S2C/Table S4 [32–40 hpi]).

Notably, we observed far more differences in *Pf*Coronin-associated proteins between WT and mutant lines in ring-stage parasites than at later stages. In both developmental stages, the major interacting partner of *Pf*Coronin was *Pf*Actin (PF3D7_1246200; Figure 1A-B), consistent with the known role of coronins in actin regulation^19,2014,23,24^. However, in ring stage, this interaction was reduced threefold in mutant parasites (*p* = 0.0113; Figure 1A and Figure S2B; Table S3), while no significant change was detected at late-stage parasites (*p* = 0.4787; Figure 1B and Figure S2C; Table S4). These results demonstrate that *Pfcoronin* mutations specifically disrupt *Pf*Coronin–*Pf*Actin association during the ring stage, coinciding with the window of ART susceptibility.

**Figure 1.**
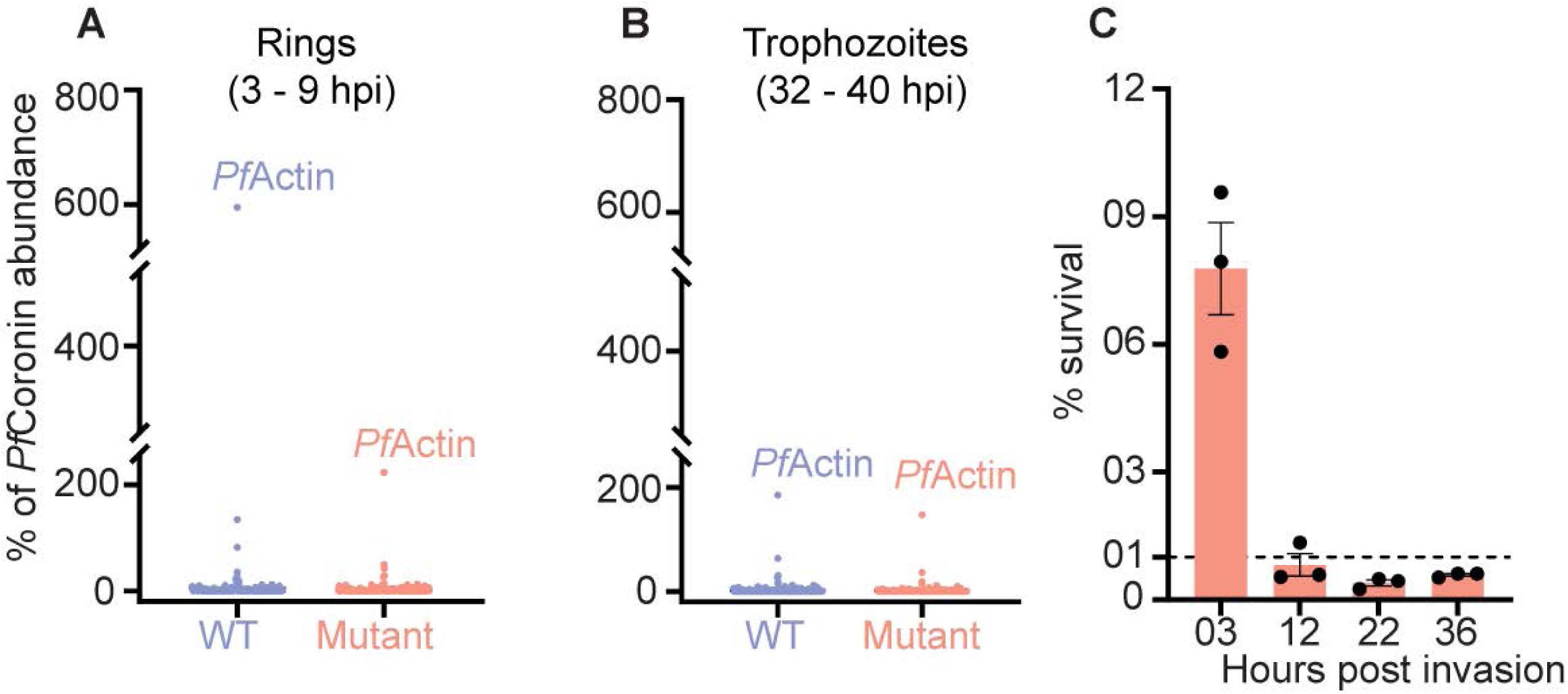
*Pfcoronin* mutations reduce *Pf*Coronin–*Pf*Actin association in ring-stage parasites. Immunoprecipitation of *Pf*Coronin-smHA and associated proteins in ring (**A**) and (**B**) late trophozoite-stage parasites (n=4 rings; n = 6 trophozoites). **C.** RSA survival (%) of *Pfcoronin* mutant line following 700 nM DHA pulse for 6-hour increments at various points in the intraerythrocytic developmental cycle.

### *Pfcoronin* mutations only confer ART resistance in rings

This molecular specificity was mirrored at the phenotypic level: comparison of *Pf*Coronin interaction partners between WT and mutant parasites showed many more differences at ring-stage than in late-stage parasites. Interestingly, this corresponds to the timing of ART clinical resistance, which manifests as delayed clearance of early ring-stage parasites. Additionally, *Pfkelch13*-mediated ART resistance in cultured parasites is only evident at the ring stage^4,6^. The ring-stage survival assay (RSA), where ring-stage parasites are exposed to physiological concentrations of dihydroartemisinin (DHA) for just 6 hours, has become the standard assay for *in vitro* ART resistance, with greater than 1% survival indicating ART resistance^4^.

To assess whether *Pfcoronin*-mediated ART resistance is stage-specific, we treated parasites with 700 nM DHA for 6-hour pulses at various timepoints across development. *Pfcoronin* mutant line (previously reported DHA-selected line carrying the R100K/E107V mutations; see Table S5)^12^ showed no resistance phenotype beyond 9 hpi (Figure 1C). Thus, like *Pfkelch13* mutants, *Pfcoronin*-mediated ART resistance is strictly ring-stage specific. This precise temporal restriction implicates *Pfcoronin* as a central determinant of parasite susceptibility to ART during early intraerythrocytic development, matching the timing of clinical resistance observed in patients.

### *Pf*Coronin WD40 propeller mutations reduce its abundance in ring-stage parasites

Mutations in WD40 β-propeller domains frequently impair protein stability and reduce steady-state abundance across diverse systems, including *Pfkelch13* mutations in *P. falciparum*^25–30^. *Pf*Coronin also contains a WD40 propeller domain, and resistance-associated mutations R100K/E107V lie within this conserved structural fold. Immunoblot analysis confirmed a marked reduction in *Pf*Coronin abundance in the mutant line at the ring stage (3–9 hpi) (Figure 2A-B; see Figure S3 for full western blot images and additional examples). This stage-specific protein loss was robust across both BiP and aldolase loading controls (*p* = 0.0054; Figure 2A-B), and levels of BiP, aldolase, and GAPDH remained unchanged between genotypes (Figure 2C). No significant difference in *Pf*Coronin abundance was observed in late trophozoite-stage parasites (Figure 2D). These data indicate that mutations induce a stage-specific reduction in *Pf*Coronin protein abundance without broadly affecting the parasite proteome. Collectively, these findings suggest that destabilization of *Pf*Coronin during the early ring stage may impair its functional capacity, potentially contributing to altered cytoskeletal regulation and parasite susceptibility to ART.

**Figure 2.**
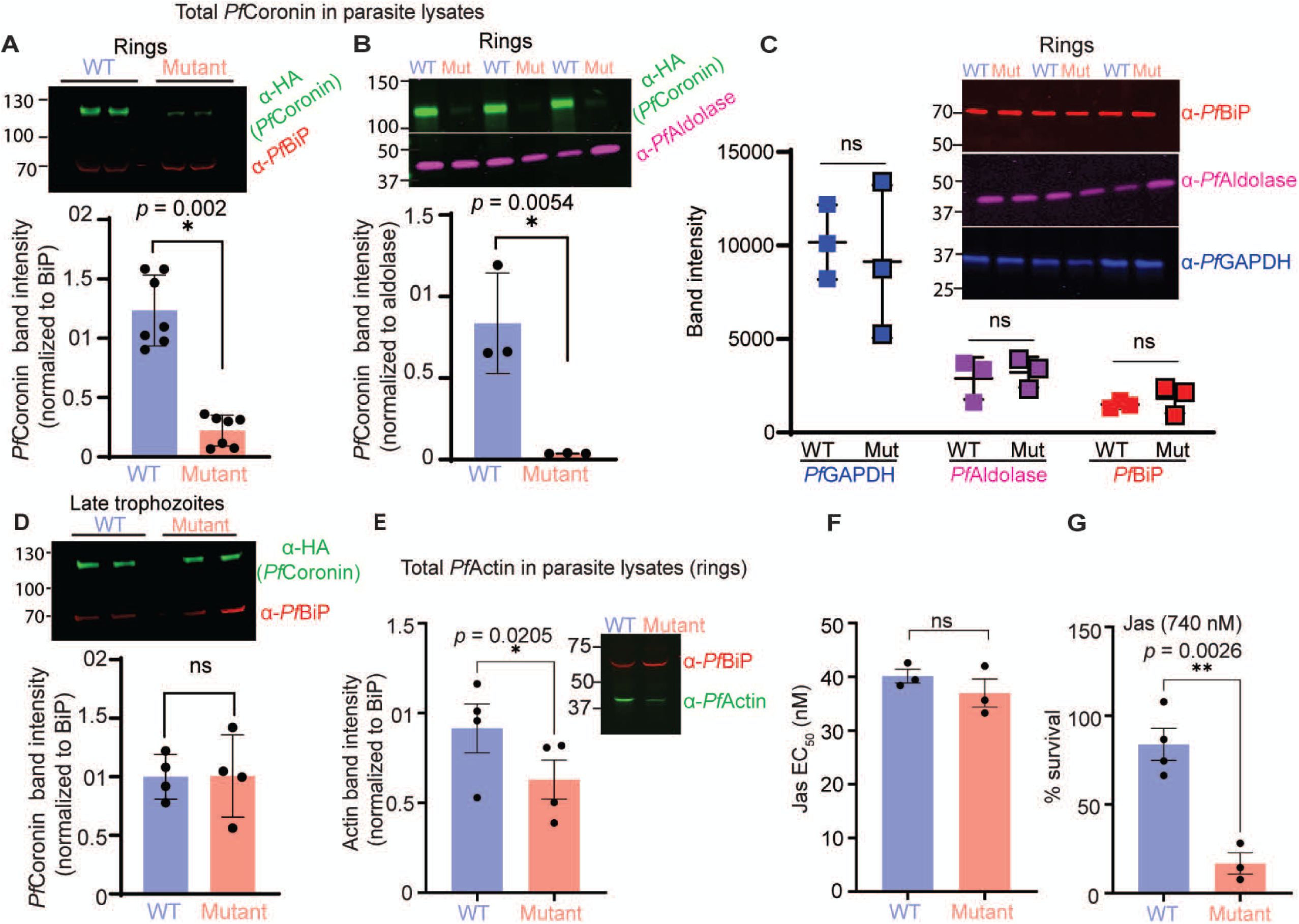
***Pfcoronin* mutations destabilize the protein in rings and disrupt *Pf*Actin homeostasis A & B.** Western blot analysis comparing total *Pf*Coronin levels in lysates from both WT and mutant ring-stage parasites. Representative blots are shown. Blots were probed with anti-HA antibodies (*Pf*Coronin; mouse monoclonal; green) and loading control antibodies: anti-*Pf*BiP (rabbit polyclonal; red; **A**) and anti-*Pf*Aldolase (rabbit polyclonal; magenta; **B**) for rings. Bar graph shows densitometric quantification of *Pf*Coronin, normalized to either *Pf*BiP (**A**) or *Pf*Aldolase (**B**). **C**. Total *Pf*BiP, *Pf*Aldolase, and *Pf*GAPDH protein levels were compared between WT and mutant (Mut) ring-stage parasites. **D**. *Pf*Coronin protein levels between WT and mutant parasites in late trophozoites. **E.** Total *Pf*Actin levels in lysates from ring-stage parasites expressing mutant *Pf*Coronin were compared by western blot to lysates from parasites expressing WT *Pf*Coronin (representative blot shown). Blots were probed with anti-actin (mouse monoclonal; green) and anti-*Pf*BiP (rabbit polyclonal; red, loading control) antibodies. The bar graph shows densitometric quantification of *Pf*Actin, normalized to *Pf*BiP. **F.** E_50_ values of JAS and (**G**) percent survival of WT and mutant parasites treated for 6 hrs, starting at 3 hpi, with 740 nM JAS.

### *Pfcoronin* mutations reduce *Pf*Actin protein levels in rings

Since we observed reduced interactions of *Pf*Actin–*Pf*Coronin in mutant ring-stage parasites, we were interested in whether these changes represented global differences in *Pf*Actin protein levels. We performed western blot analysis on ring-stage parasite lysates to assess whether *Pf*Actin abundance differs between WT and mutant parasites. We found that mutant parasites contained 32.3 ± 6 % (mean ± SEM) less *Pf*Actin, compared to WT parasites in rings (*p* = 0.0205; Figure 2E; see Figure S4A for full western blot images and additional examples). This decrease is consistent with the observed loss of *Pf*Coronin–*Pf*Actin interaction and suggests a possible reduction in *Pf*Actin stability or altered turnover in the mutant background. We did not observe changes in *Pf*Actin protein levels between WT and mutant in late-stage parasites (Figure S3B). Our results show that *Pfcoronin* mutations selectively destabilize *Pf*Actin during the ring stage—highlighting the importance of precise cytoskeletal regulation in defining drug response.

### *Pfcoronin* mutants sensitize *Pf*Actin to jasplakinolide

Our data revealed that mutant parasites have changes in *Pf*Actin abundance. We therefore hypothesized that *Pfcoronin* mutations lead to defects or dysregulation of *Pf*Actin dynamics. We reasoned that these changes might alter sensitivity to jasplakinolide (JAS), which stabilizes *Pf*Actin filaments and promotes polymerization, ultimately disrupting normal *Pf*Actin dynamics. We tested this hypothesis using WT and mutant parasites (DHA selected line; Table S5)^12^. We first estimated sensitivity to JAS using a standard 72 h growth assay and found no difference between WT (EC_50_ = 40.1 ± 1.3 nM) and mutant parasites (EC_50_ = 37 ± 2.6 nM; Figure 2F and Figure S5). To assess possible differences in ring-stage parasites, we performed RSA-type assays on WT and mutant parasites. Parasites were treated for 6 hrs, starting at 3 hpi, with 740 nM JAS (∼20X EC_50_). JAS had a modest effect on WT parasite survival (84 ± 9% survival), but significantly reduced the survival of mutant parasites (17 ± 6% survival) (Figure 2G). Thus, *Pfcoronin* mutations sensitize parasites to *Pf*Actin disruptions, suggesting some level of underlying *Pf*Actin dysfunction or dysregulation, particularly in early ring-stage parasites.

### *Pfcoronin* mutations disrupt membrane localization in ring-stage

To gain further insights into *Pf*Coronin localization, we performed ultrastructure expansion microscopy (U-ExM)^31–33^, assessing *Pf*Coronin localization. In WT ring-stage parasites, *Pf*Coronin was found at the membranes, likely representing the parasite plasma membrane (PPM) (Figure 3). By contrast, in mutant parasites, *Pf*Coronin displayed disorganized puncta at the ring stage, consistent with the overall ring-stage specific phenotype of *Pf*Coronin.

**Figure 3.**
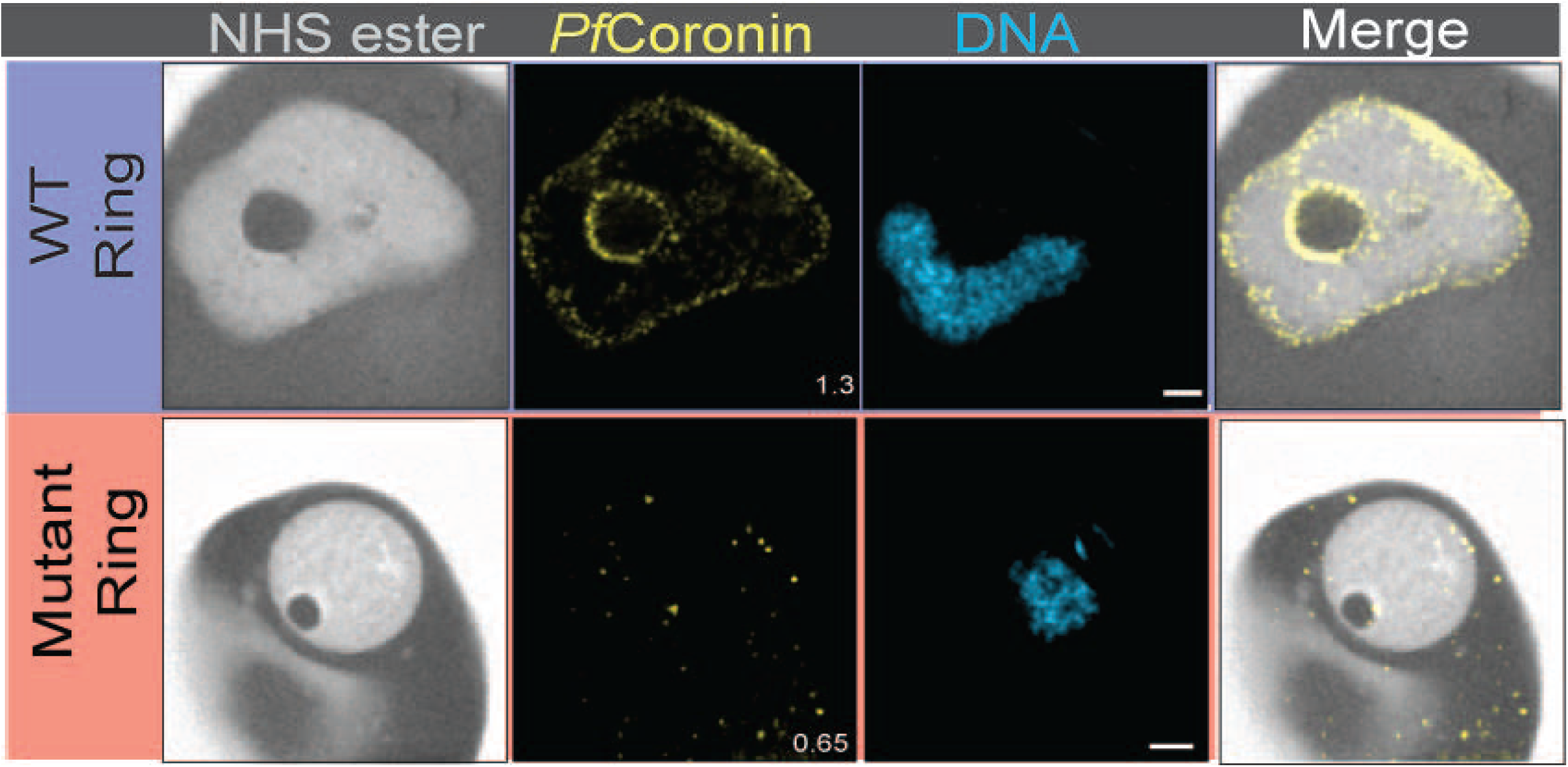
***Pfcoronin* mutations disrupt *Pf*Coronin localization in ring-stage parasites.** Super-resolution imaging using U-ExM on a Zeiss LSM900 with Airyscan2 was used to visualize smHA-tagged *Pf*Coronin WT and *Pf*Coronin mutant in ring-stage parasites. Alexa Fluor 405–conjugated NHS ester labels protein density (gray), anti-HA labels *Pf*Coronin (yellow), and Sytox Deep Red stains DNA (cyan). Images are shown as maximum-intensity projections; numbers on images indicate the Z-axis thickness of the projection (µm). Scale bars, 2 µm.

### *Pf*Coronin marks vesicle trafficking intermediates and the digestive vacuole (DV) membrane

To gain further insight into the functional roles of *Pf*Coronin in parasite cellular architecture, we examined its subcellular localization in late-stage parasites. In WT parasites, *Pf*Coronin was enriched at the DV membrane (Figure 4A) and was frequently observed on chains of small and large vacuoles (Figure S6), possibly representing vesicle trafficking intermediates en route to the DV. Co-staining with anti-aldolase confirmed that these vacuoles are distinct from the parasite cytoplasm (Figure S6). In contrast, *Pfcoronin* mutants exhibited disrupted localization of *Pf*Coronin to both the DV membrane and associated vacuolar structures (Figure 4A). These localization patterns were corroborated by conventional immunofluorescence assays (IFA), further supporting the observations obtained by U-ExM (Figure S7) U-ExM further enabled visualization of small organelles such as the cytostome, where *Pf*Kelch13 localizes to the neck region; *Pf*Kelch13 has also been implicated in cytostome-mediated hemoglobin uptake^6,32,34^. The cytostome is a double-membrane structure comprised of both the PPM and the parasite vacuolar membrane (PVM), forming distinct neck and body regions^6,32,34^. We occasionally observed *Pf*Coronin-coated vacuoles in proximity to the cytostome (Figure 4A). Although direct association between *Pf*Coronin and the cytostomal body was not detected, given *Pf*Coronin’s enrichment at the PPM and the double-membrane architecture of the cytostome, potential association of *Pf*Coronin with the cytostome neck cannot be ruled out.

**Figure 4.**
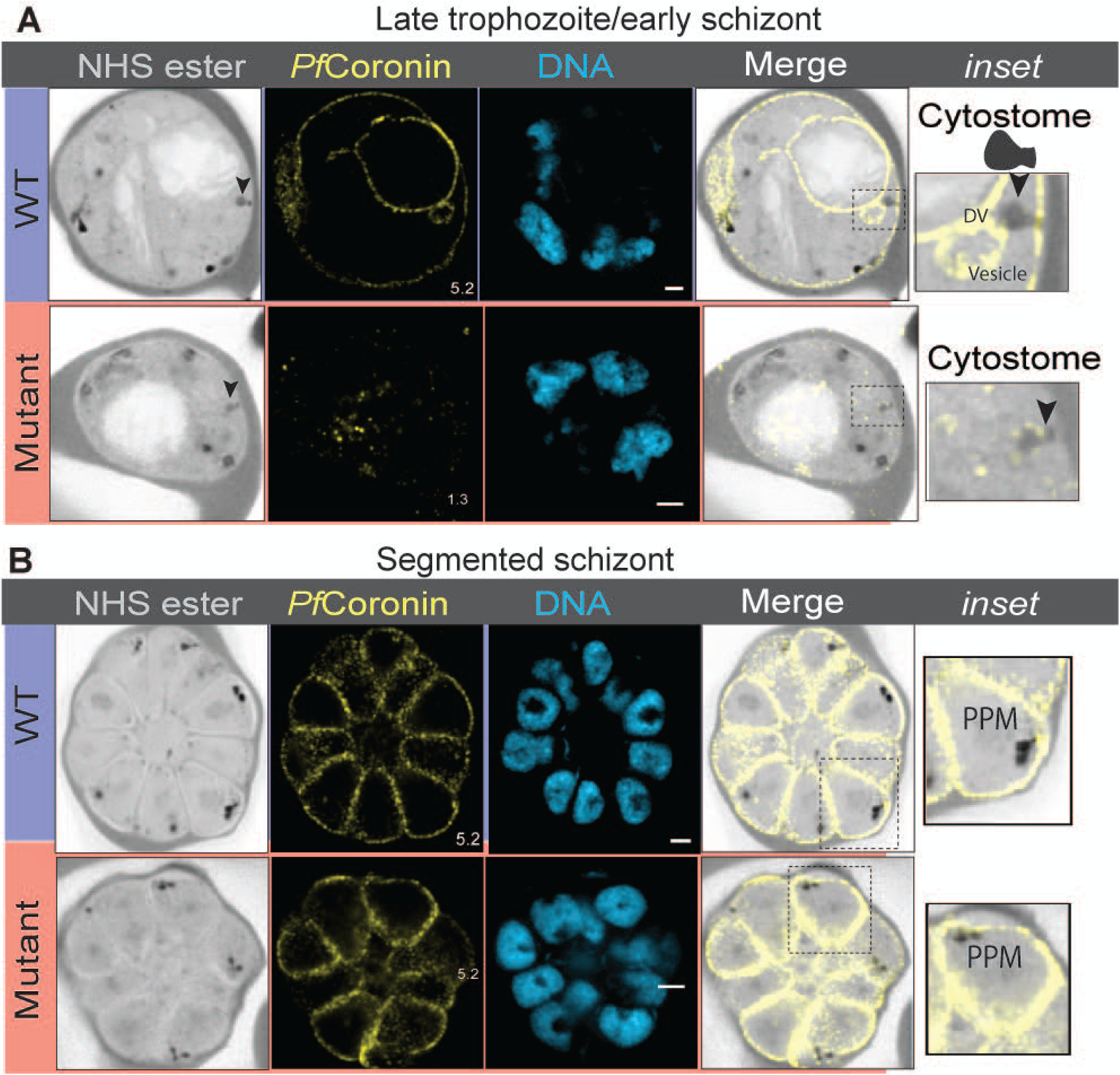
***Pf*Coronin localization to the DV and vesicles is disrupted by mutation and restored only in segmented schizonts.** Super-resolution imaging using U-ExM on a Zeiss LSM900 with Airyscan2 was used to visualize smHA-tagged *Pf*Coronin (WT and mutant) in late-stage parasites. **A.** Comparison of *Pf*Coronin localization in WT and mutant parasites at the late trophozoite/early schizont stage. A schematic of the cytostome is shown in inset, with an arrowhead indicating its position. **B.** Comparison of *Pf*Coronin localization in WT and mutant parasites at the segmented schizont stage. Alexa Fluor 405–conjugated NHS ester labels protein density (gray), anti-HA labels PfCoronin (yellow), and Sytox Deep Red stains DNA (cyan). Images are shown as maximum-intensity projections; numbers indicate the Z-axis thickness of each projection (µm). Scale bars, 2 µm.

### *Pf*Coronin mutant localization restores to normal in schizonts

U-ExM during late schizogony showed that *Pf*Coronin localization in mutant parasites returns to a WT-like pattern; the earlier mislocalization is resolved in segmented schizonts (Figure 4B). At this stage, both *Pf*Coronin WT and mutant are localized to the periphery of individual merozoites, consistent with previous reports^21^.

We next attempted to assess colocalization of *Pf*Coronin with its binding partner, *Pf*Actin. *Pf*Actin was broadly distributed throughout the cytoplasm and frequently detected near *Pf*Coronin at the periphery, consistent with previous studies^21^. No substantial differences in *Pf*Actin distribution were observed between WT and mutant lines (Figure S8).

To further investigate whether altered *Pf*Coronin localization impacts parasite development or intraerythrocytic progression, we closely monitored both WT and mutant parasites throughout intraerythrocytic cycle. Despite pronounced stage-specific mislocalization and depletion of *Pf*Coronin at the ring stage, both mutant and WT parasites displayed indistinguishable morphology, developmental timing, and size throughout development (see representative images in Figure S9). While our previous work established that *Pf*Coronin mutations do not impair parasite growth over multiple cycles^13^, these data further demonstrate that there is no detectable short-term developmental arrest or delay within a single asexual cycle, suggesting that the canonical function of *Pf*Coronin is not required for asexual stage progression *in vitro*.

Collectively, these results indicate that *Pf*Coronin is a temporally regulated, non-essential protein whose stage-specific destabilization by resistance-associated mutations leads to distinct cellular and molecular changes without broadly impairing parasite development.

### Mutations in *Pfcoronin* reduce hemoglobin uptake

Prior studies demonstrated that the uptake of red blood cell (RBC) contents is decreased in ART-resistant *Pfkelch13* mutants^5,6^. Given that ART activation depends on free heme produced via endocytic uptake of host hemoglobin, *Pf*Coronin’s association with vesicular structures led us to hypothesize that it may influence the endocytic pathway and thereby modulate ART susceptibility, particularly during the ring stage. To test this hypothesis, we performed a previously published microscopy-based endocytosis assay, using fluorescent-dextran as a tracer for uptake of host cell contents^5^ (Figure S10). Since *Pfkelch13* mutants (with C580Y mutation) had previously been assessed in the same assay^6^, we used *Pfkelch13* mutants (C580Y)^13^ to validate the assay. We measured a significant 40.3 ± 16.6% (mean ± SD) reduction in fluorescent-dextran uptake in *Pfkelch13* mutant (C580Y) parasites compared to the WT parent (unpaired Mann-Whitney test; *p* < 0.0001; Figure S11A; see Figure S11B-D for individual bioreps). This figure closely matches the 37.47% reduction previously reported in *Pfkelch13* mutants (C580Y)^5^. We then assessed fluorescent-dextran uptake in *Pfcoronin* mutants. Interestingly, we observed a 38.8 ± 14.4% (mean ± SD) reduction in fluorescent-dextran uptake in *Pfcoronin* mutants compared to *Pf*Coronin WT parasites (unpaired Mann-Whitney test; *p* < 0.0001; Figure 5; see Figure S12 for individual bioreps).

**Figure 5.**
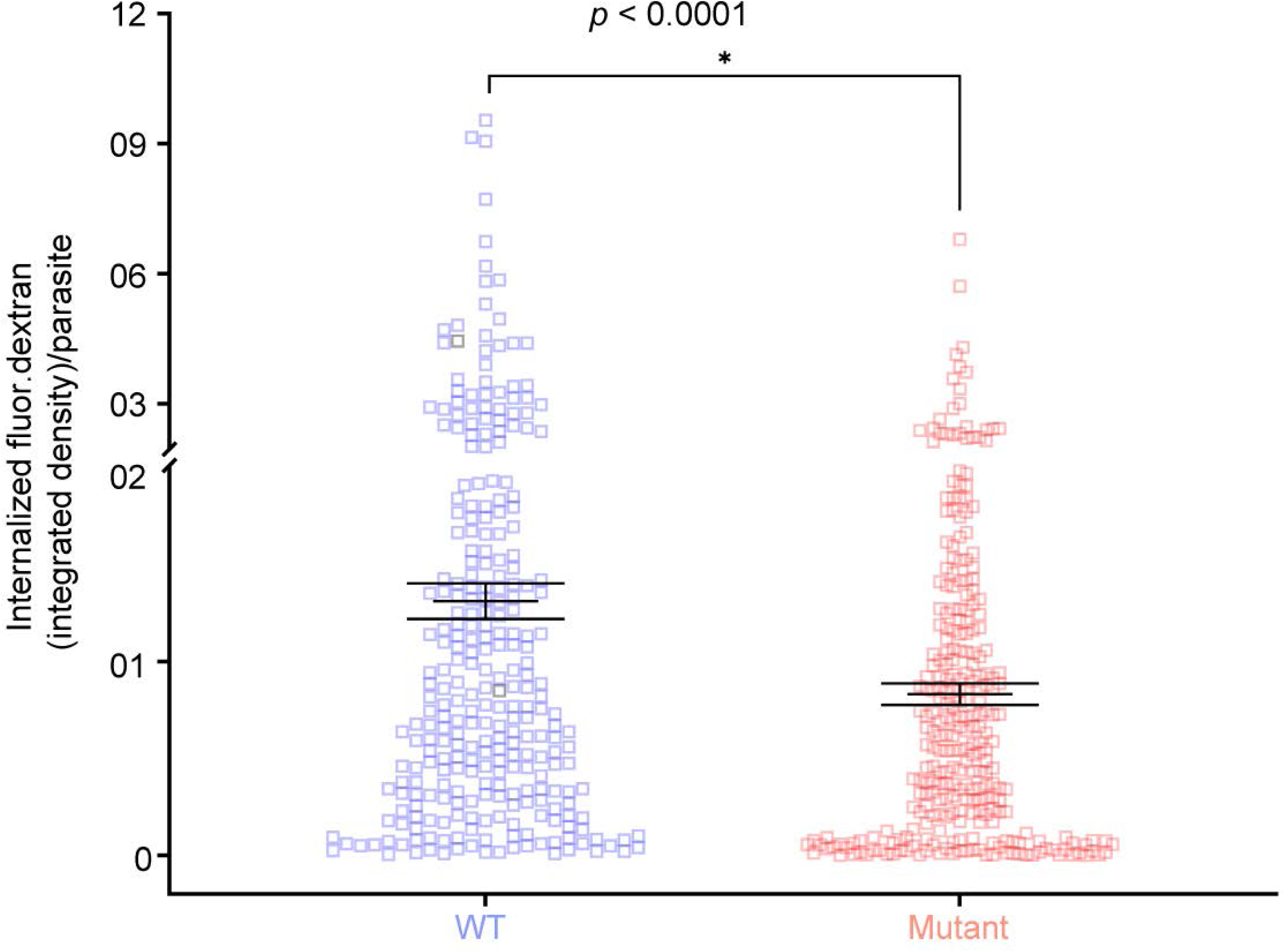
***Pf*Coronin mutants decrease host cell content uptake** Comparison of internalized fluorescent dextran (fluor.dextran) in parasites expressing *Pf*Coronin WT versus *Pf*Coronin mutant, measured by integrated density (vesicle size × intensity). WT *Pf*Coronin: n = 286; mutant *Pf*Coronin: n = 286. Each data point represents fluorescent dextran uptake by a single parasite. *p*-values (unpaired, nonparametric Mann-Whitney test) are indicated. Data were pooled from the individual biological replicates shown in Figure S12. Non-smHA tagged parasites, *Pf*Coronin WT, *Pf*Coronin mutant, and *Pf*Kelch13 C580Y (all in the Pikine background; see Figure S10 for *Pf*Kelch13) were included in these analyses.

Together, these results reveal that the non-essential cytoskeletal regulator *Pf*Coronin, while not required for asexual parasite growth, contributes to efficient hemoglobin uptake and optimal ART susceptibility at the clinically relevant ring stage. Mutations in *Pfcoronin* mimic the effects of *Pfkelch13* variants by restricting endocytic access to host hemoglobin, thereby limiting heme-dependent drug activation. The distinct localization of *Pf*Coronin points to engagement with parallel or downstream endocytic pathways, highlighting how non-essential, stage-specific parasite factors can be co-opted to diversify cellular strategies for therapeutic evasion and drug resistance.

## Discussion

In this study, we show that the non-essential cytoskeletal regulator *Pf*Coronin has a stage-specific role in endocytic trafficking and ART susceptibility in *P. falciparum*. Resistance-associated *Pfcoronin* mutations (R100K/E107V) confer ART resistance only during the early ring stage. These mutations perturb *Pf*Coronin abundance, subcellular localization, and its association with *Pf*Actin only at the ring stage, while later developmental stages are unaffected. Accompanying phenotypes—including decreased *Pf*Coronin–*Pf*Actin interaction, reduced *Pf*Actin levels, and hypersensitivity to the actin-stabilizer JAS—are likewise limited to early rings. U-ExM analysis further reveals that WT *Pf*Coronin is distributed at the PPM, DV, and vesicular compartments, but this organization is lost in mutant parasites, and is only restored in late schizogony. Functionally, *Pfcoronin* mutations reduce host hemoglobin uptake in rings—a phenotype also observed in *PfKkelch13*-mediated ART resistance^5^. Together, these findings establish that non-essential, stage-specific cytoskeletal regulators can influence endocytic access to host resources and thus modulate drug response in malaria parasites.

Distinctive features of *P. falciparum* endocytosis offer opportunities for selective therapeutic intervention. *Pf*Kelch13 localizes to the cytostome neck, and ART resistance–associated mutations in *Pf*kelch13 and interacting partners reduce hemoglobin uptake and digestion^4,35,36^. Unlike mammalian cells, where endocytosis is clathrin-dependent, the parasite cytostome was previously thought to lack clathrin but incorporates AP2-μ, highlighting evolutionary adaptations^5^. However, recent evidence^37^ implicating clathrin and AP1 in cytostome organization suggests this model may need revision. Given these differences, understanding how additional regulators contribute to endocytosis and drug resistance is critical.

Our localization studies show that *Pf*Coronin supports hemoglobin uptake from sites beyond the cytostome neck—specifically, at the PPM, DV, and vesicular compartments—suggesting a broader regulatory function. Although *Pf*Coronin lacks canonical lipidation motifs or transmembrane domains; mammalian coronins are known to associate with membranes through interactions with phosphatidylinositol 4,5-bisphosphate (PI(4,5)P2)^38^. *Pf*Coronin localizes adjacent to the cytostome neck on the PPM, without overlapping the cytostomal body, suggesting it regulates *Pf*Actin dynamics downstream of vesicle formation. This likely coordinates actin-dependent vesicle transport and maturation during hemoglobin uptake, though alternative uptake pathways cannot be excluded. This role aligns with conserved coronin functions in other eukaryotes^39,40^. No direct physical association between *Pf*Coronin and *Pf*Kelch13 was detected here or in prior studies^5^, supporting a model in which these proteins contribute to separate, yet potentially coordinated, steps of endocytosis. Previous genetic analysis^13^ indicates possible functional crosstalk or compensation between *Pfcoronin* and *Pfkelch13*, even though the molecular basis remains to be defined.

Actin cytoskeleton dynamics are central to endocytic processes in eukaryotes, facilitating vesicle trafficking and efficient nutrient acquisition^41^. In *P. falciparum*, stabilization of *Pf*Actin filaments disrupts cytostome architecture and hampers hemoglobin delivery to the DV^42^. We find that *Pfcoronin* mutations alter both *Pf*Actin abundance and its functional regulation, as evidenced by increased JAS sensitivity in ring stages. Apicomplexan actins are short and highly dynamic^43^, and as widely recognized in the field, conventional antibodies—including those used in this study—cannot reliably distinguish monomeric from filamentous *Pf*Actin, limiting their ability to capture these transient structures. Chromobody-based reporters provide live-cell, high-resolution visualization of *Pf*Actin filaments in their native state^44^,and employing this approach in future studies will be critical to define how *Pf*Coronin interfaces with the *Pf*Actin network and to determine how *Pfcoronin* mutations perturb this interplay during parasite development. While the core trafficking machinery in *P. falciparum* is less understood than in model systems^5,45–48^, the parasite encodes a limited but specialized set of *Pf*Actin regulators, among which *Pf*Coronin is notable for its non-essential but regulatory role^49–52^. Beyond *Pf*Kelch13 and partners, relatively few proteins—including phosphoinositide-binding protein PX1^53^, VPS45^46^, and the host enzyme peroxiredoxin 6^47^—have been linked to endocytic trafficking. Our data highlight *Pf*Coronin as an additional candidate in this restricted but functionally vital network.

Our results show that resistance-associated mutations in *Pfcoronin* selectively disrupt its subcellular localization and reduce protein abundance during the ring stage—the window of maximal ART susceptibility. This effect is absent in segmented schizonts, where *Pf*Coronin localization patterns are similar in both genotypes, but differences emerge at earlier stages. In trophozoites, while overall *Pf*Coronin abundance is comparable, mutants lack the spatial membrane association characteristic of WT parasites. This reflects disrupted recruitment or retention at key subcellular sites, likely compromising its role in trafficking or cytoskeletal organization. These findings support the idea that *Pf*Coronin undergoes stage-specific regulation, potentially involving turnover, trafficking, or interaction with short-lived partners. Loss of membrane localization in rings and trophozoites— developmental windows of active hemoglobin uptake—indicates a disruption in the coordination between membrane and cytoskeletal systems. In contrast, normalization of *Pf*Coronin localization in schizonts may reflect a reduced requirement for *Pf*Coronin-mediated processes or compensation by other cytoskeletal components at this late stage. Thus, the restoration of *Pf*Coronin localization in schizonts remains intriguing, and may reflect stage-specific regulatory mechanisms independent of endocytic activity.

The mechanism underlying the stage-specific disruption of mutant *Pf*Coronin localization remains unclear. One possibility is that *Pf*Coronin requires transient interactors or stabilizing factors—such as membrane adaptors, lipid domains, or actin-associated proteins—that are specifically expressed or active during the ring and trophozoite stages but are downregulated or absent in segmented schizonts. Alternatively, the mutant protein’s altered conformation may impair its ability to engage with such partners under the dynamic trafficking conditions of early stages, but this defect becomes masked or inconsequential when trafficking ceases at schizogony. Our current interactome approaches, which emphasize stable and abundant binding partners, likely miss these transient or low-abundance interactors. To overcome these limitations and fully define *Pf*Coronin’s functional network across the parasite’s developmental cycle, future studies should employ stage-specific proximity labeling. Such approaches are poised to reveal how resistance-associated mutations modulate *Pf*Coronin’s connectivity with the *Pf*Actin network and other trafficking regulators, offering a high-resolution view of how this non-essential cytoskeletal protein mediates drug susceptibility and ring-stage biology.

Mechanistically, *Pf*Coronin’s regulatory role in vesicular trafficking manifests exclusively in ring stages. Although dispensable for *in vitro* growth, altered *Pf*Coronin impairs endocytosis and drug responsiveness when present in its mutant form. This highlights how “non-essential” proteins can nevertheless provide critical, stage-restricted functions emphasizing the importance of reassessing gene essentiality in the context of parasite development and drug response networks. Clinically, these findings are timely. Recent identification of *Pfcoronin* mutations, including R100K, in African isolates—some linked to delayed parasite clearance and reduced ACT efficacy^8–11^—emphasizes the need for systematic surveillance of ART resistance markers beyond canonical pathways. Our data underscore that adaptations in cytoskeletal and endocytic mechanisms help drive parasite resistance evolution, and that conventional notions of gene essentiality may overlook stage- or pathway-specific vulnerabilities vital for parasite fitness and drug sensitivity.

In summary, *Pf*Coronin exemplifies how non-essential, stage-specific regulators orchestrate the interface between host resource acquisition and antimalarial drug susceptibility. *Pf*Coronin acts as a molecular organizer whose dysfunction—rather than simple absence—undermines hemoglobin uptake and drug response. This highlights the complexity of ring-stage biology: a protein that is dispensable for overall survival can nonetheless be important in specific cellular functions and drug sensitivity, likely due to its role within a larger, stage-specific protein interaction network. As ART remains central to global malaria treatment, ongoing mechanistic studies and field genetic surveillance of *Pfcoronin*—and other adaptive regulators—will be essential for effective malaria control and strategies to anticipate evolving drug resistance.

## Resource Availability

### Lead Contact

Further information and requests for resources and reagents should be directed to, and will be fulfilled by, the lead contact: Dyann F. Wirth.

## Materials availability

All cell lines and plasmids in this study are available from the lead contact upon completion of a materials transfer agreement.

## Data and code availability

Data availability:

The data that support the findings of this study are included in the supplementary materials and source data are provided with this manuscript. Any additional data related to this study can be requested from the corresponding author.

## Code availability

This paper does not report codes.

## Acknowledgments

Funding: NIH (5R01AI099105-07) to DFW and DLH and NIH (R01 AI145941) to JDD. This work was supported in part by the Undergraduate Research Opportunity Program (UROP) at Indiana University (SA).

## Author contributions

Conceptualization: I.U., J.D.D., D.L.H., S.K.V., S.B., S.A., and D.F.W.; Methodology: I.U., M.A.F., A.Y.B., E.H., B.C.W., M.K., S.H.S., A.I.S., K.L.S., M.C.M., S.A.; Data analyses: I.U., B.C.W., M.K., S.A.; Writing - original draft: I.U.; writing - review & editing: I.U., B.C.W., J.D.D., D.L.H., S.K.V., S.B., S.A., and D.F.W.; Funding acquisition: D.F.W., and D.L.H., Supervision: D.F.W.

## Declaration of generative AI and AI assisted technologies in the writing process

The authors wrote and edited the manuscript and take full responsibility for the content of the published article.

## Supplemental information

Document S1. Figures S1–S12 and Table S5.

Table S1. Ring-stage *Pf*Coronin-associated proteins.

Table S2. Late trophozoite-stage *Pf*Coronin-associated proteins.

Table S3. High-confidence interaction partners of *Pf*Coronin in ring-stage parasites, related to Figure 1A and Figure S2B.

Table S4. High-confidence interaction partners of *Pf*Coronin in late trophozoite-stage parasites, related to Figure 1B and Figure S2C.

## STAR Methods

### Parasite culture

*P. falciparum* culture-adapted field isolates, Pikine (SenP019.04), Pikine_R (DHA selected line^12^), CRISPR-engineered Pikine *Pfcoronin^R100K/E107V^* mutant (CRISPR-edited), *Pf*kelch13^C580Y^ mutant (CRISPR-edited)^12,13^, spaghetti monster HA tag (smHA)-tagged Pikine *Pfcoronin^WT^-smHA*, and Pikine *Pfcoronin^R100K/E107V^-smHA* mutant lines were maintained in complete RPMI 1640 medium (Gibco) supplemented with 10% O^+^ serum at 4% hematocrit (unless specified otherwise) in an atmosphere of 1% O2, 5% CO2, and 94% N2 at 37 °C^12,13^. All smHA-tagged lines were additionally cultured with 5nM WR99210 (Jacobus Pharmaceutical). Staging and parasitemia were assessed by light microscopy of Giemsa-stained thin blood smears^54,55^. The parasites were synchronized using sequential sorbitol lysis treatment ^56^. All cell lines utilized in this study are described in detail in **Table S5**.

### Generation of smHA-tagged parasites and transfection

*Pfcoronin* (PF3D7_1251200) was tagged with smHA using CRISPR/Cas9. The CRISPR guide sequence (selected using benchling.com) was 5’-AATGTGTAAAAGTACAGCAA-3’, which matches a site near the C-terminus of the *Pf*Coronin coding sequence. A double-stranded DNA segment with “sticky ends” was generated by annealing oligonucleotides 5’-ATTGAATGTGTAAAAGTACAGCAA-3’ and 5’-AAACTTGCTGTACTTTTACACATT-3’. This segment was ligated within the U6 cassette of pBAM203^57^ (pre-digested with BbsI; kind gift of Dr. Jeffrey Dvorin, Harvard Medical School), proximal to the invariable part of the chimeric sgRNA scaffold, to generate pUF1-Cas9-hDHODH-Coronin. The donor/homologous repair plasmid for smHA-tagging *Pf*Coronin was derived from pSAB55^32^ (kind gift of Dr. Jeffrey Dvorin, Harvard Medical School), which contains coding sequence for spaghetti monster-HA tag (smHA), followed by the *Pf*HSP86 3’ UTR. Additionally, the plasmid includes a cassette for expression of human dihydrofolate reductase (hDHFR), allowing selection using WR99210. A 5’ homology arm (plus shield mutations) was amplified from the end (C-terminus) of the *Pf*Coronin coding sequence using primers: 5’-CAATTGCCATGGGACAGTTTACCAAAAAATTTACCTTC-3’ and 5’-ATCTCGAGTAAAACGGTTGCAGTACTTTTACACATTTTCATGTCTAG-3’; the second primer includes shield mutations to protect from cutting by Cas9. This segment was inserted into the NcoI/XhoI sites of pSAB55. A 3’ homology arm was amplified from the *Pf*Coronin 3’ UTR using 5’-CGTCAGGATCATCGCGGCCGCGGCACATCAGCGATCAC-3’ and 5’-CATGGCAATTGAGATCTCCGAAAAATTTGTATGATATTGAAAAG-3’; this segment was inserted into the NotI/BglII sites of the plasmid using Gibson cloning, generating pSAB55-Coronin-smHA. Prior to transfection, the complete pSAB55-Coronin-smHA plasmid was linearized by digestion with NcoI. Plasmids (50 µg each of pUF1-Cas9-hDHODH-Coronin and linearized pSAB55-Coronin-smHA) were transfected into Pikine parasites carrying either *Pf*Coronin^WT^ (WT) or *Pf*Coronin^R100K/E107V^, as reported previously^13^. Transfections were performed as described previously^13,58^. Transfected parasites were allowed to recover overnight before the addition of 5 nM WR99210 (selection agent for pSAB55-Coronin-smHA) and 500 nM DSM1 (selection agent for pUF1-Cas9-hDHODH-Coronin). Drug selection with DSM1 was continued for 3 days. Parasites grew up around 3 weeks and were cloned by limiting dilutions prior to studies. Confirmation of the smHA-tagged parasites is detailed in Figure S1.

### Ring stage survival assay (RSA)

RSA was performed as described previously^12,13,59,60^. *P. falciparum* cultures at the late segmented schizont stage were purified using the Percoll gradient and allowed to reinvade for 3 hours before subjecting them to synchronization. Highly synchronized (0-3 hpi) parasites were pulsed with 700 nM DHA (Sigma-Aldrich) or 740 nM jasplakinolide (ref no, 420127, Sigma-Aldrich), or DMSO control. The pulse was followed by drug washout and further incubation at 37^°^C for 66 hours, and parasitemia was assessed by light microscopy of Giemsa-stained thin blood smears^12,13^. RSA survival rates in percentage were estimated by comparing the survival of viable parasites to the untreated DMSO control for the same parasite line. Smears were blinded, and 10,000 RBCs were counted for each condition. Unless otherwise specified, all experiments used at least three independent biological replicates and two technical replicates.

### Concentration–response assays

All EC_50_s were estimated using the standard 72 hour Malaria SYBR green I fluorescence assay^61,62^. Ring-stage parasites (40 μL, 1% hematocrit and 1% starting parasitemia, n = 3) were cultured for 72 hours at 37°C in a 384-well black clear-bottom plates containing 12-point serial dilutions of test compounds, 10 μM DHA kill-control and no drug growth-control. Lysis buffer containing 16% w/v saponin, 1.6% Triton X-100, 5 mM EDTA, 20 mM Tris-HCl, pH 7.4) and 1x SYBR Green I fluorescent dye (CAT. No. S7563, Invitrogen) was added, and the plates were incubated at room temperature in the dark for at least 6 hours prior to reading. The fluorescent signal was measured using SpectraMax M5 (Molecular Devices, Sunnyvale, CA) plate reader (excitation at 494 nm, emission at 530 nm). Experiments were carried out as three technical triplicates on the same plate, with three independent biological repeats of each plate performed. Controls in each biological replicate consisted of parasite culture with no drug added (100%) or 10 μM DHA kill-control (0%). The mean and SD of fluorescence data from three independent biological repeats were expressed as a proportion of the untreated control (100%) and calculated as follows: 100 × [μ(S) − μ(−)/μ(+) − μ(−)], where μ(S), μ(+) and μ(−) represent the means for the sample in question and 100% and 0% controls, respectively as previously described^54,55,63^. The percentage growth was plotted against log_10_-transformed drug concentration and EC_50_s were estimated using a non-linear regression (log(inhibitor) vs response-variable slope with four parameters) in GraphPad Prism (GraphPad Software, Inc., San Diego, CA, USA). Values from the three biological replicates were used to calculate the mean EC_50_ values ± SE shown.

### Co-Immunoprecipitation

Tightly synchronized *Plasmodium falciparum* parasites were obtained using 5% D-sorbitol treatment followed by Percoll density centrifugation. For each Co-IP experiment, approximately 4 × 10⁹ parasites were harvested at either 32–40 hpi (late stages) or 3–9 hpi (ring stages). Infected red blood cells (RBCs) were washed with PBS and lysed using 0.05% saponin in PBS in the presence of EDTA-free complete mini protease inhibitors (Cat. No. 4693159001, Sigma). The released parasites were washed thoroughly with PBS supplemented with protease inhibitors to remove residual RBC components. Parasite pellets were lysed in RIPA buffer (50 mM Tris-HCl, pH 7.5, 150 mM NaCl, 1% NP-40, 0.5% sodium deoxycholate, 0.1% SDS, and EDTA-free complete mini protease inhibitors) by incubation on a rotator for 1 hour at room temperature. Lysates were sonicated three times at 20% amplitude for 30 seconds with 3-minute cooling intervals on ice. Insoluble material was removed by centrifugation at 12,000 × g for 30 minutes at 4 °C, and the clarified supernatant was incubated with 40 μL anti-HA magnetic beads (Cat. No. 88836, Pierce) overnight at 4 °C. Beads were then washed and submitted for on-bead digestion followed by peptide identification via LC–MS/MS, as described previously^58^.

### Immunoblotting

Synchronized parasites were isolated on ice for 10 minutes with 0.015 % saponin. Isolated parasites were washed three times with ice-cold PBS containing a protease inhibitor cocktail (Roche cOmplete mini, Sigma Cat. No. 11836153001) before lysis in 1x Laemmli Sample Buffer (Biorad Cat. No. 1610747) containing 5% beta-mercaptoethanol, followed by heating the samples at 99°C for 10 minutes. Protein samples were separated on Mini-PROTEAN® TGX™ Precast Gels (4–20% gradient, BioRad Cat. No. 4561091) in Tris-SDS-glycine buffer. Separated proteins were transferred to a nitrocellulose membrane (Life Technologies Cat. No. IB3010-31) using an iBlot system (Life Technologies). The membrane was blocked in Intercept® (TBS) Blocking Buffer (LI-COR Biosciences Cat. No. 927-60001) for 1 hour at room temperature on an orbital shaker. Membrane-bound proteins were first probed with primary antibodies [mouse anti-*Pf*Actin 5H3 (1:100; Walter and Eliza Hall Institute of Medical Research Cat. No. 26/11-5H3-1-2)^23^, mouse anti-HA (1:2000; Pierce Cat. No. 26183), , rabbit anti-aldolase (1:5000; Abcam Cat. No. ab207494), Anti-GAPDH (P. falciparum) Antibody, clone 1.4 (1:1000; Sigma Cat. NO. MABS1946) or rabbit polyclonal anti-BiP (1:5000; kindly gifted by Dr. Jeffrey Dvorin, Harvard Medical School)] in Intercept® (TBS) with 0.2% Tween 20 at 4 °C overnight. The membranes were then incubated for 45 minutes at room temperature with secondary antibodies [IRDye® 680RD goat anti-mouse (1:10,000; LI-COR Biosciences Cat No. 926-68070) and IRDye® 800CW Goat anti-Rabbit (1:10,000; LI-COR Biosciences Cat. No. 926-32211)], washed, and imaged, then analyzed using the LI-COR Odyssey CLx Imager System (LI-COR Biosciences).

### Microscopy-based endocytosis assay (ring-stage)

RBCs were suspended in a hypotonic solution and pre-loaded with fluorescent-dextran (Alexa 488; ThermoFisher Cat. No. D22910) as previously described^6,46^. After resealing red blood cells (RBCs) and trapping fluorescent dextran within, schizonts were added and allowed to re-invade by following the RSA setup. As the parasites (rings) grew within the fluorescent-dextran-containing RBCs, fluorescent-dextran was endocytosed and accumulated in the DV or other compartments. The parasites were then treated with 5% sorbitol to remove any remaining schizonts, closely mimicking the RSA setup. Parasites were washed once with RPMI medium and released from the RBCs using 10 pellet volumes of PBS containing 0.015% saponin. Following 3 washes with RPMI medium, parasite samples were fixed with pre-warmed 4% paraformaldehyde-0.1% glutaraldehyde in PBS for 20 minutes at 37°C. Afterwards, 2-5 µL samples were loaded onto clear slides, covered with a coverslip, and allowed to settle for up to 15 minutes at room temperature before imaging. DIC and AF488 channel images were captured using a 100X oil immersion objective on a Zeiss Axio Observer. AF488 channel images were acquired at 480 ms across all samples, with a laser intensity of 40%. Images were blinded prior to analysis. Analysis was conducted for regions of interest (ROIs) where an RBC ghost could be visualized around a green channel signal in the merged image. A global threshold was set on 8-bit images of the green channel across images taken at the same exposure settings. Area, mean gray value, and integrated density of the parasite-specific green-channel signal were measured using ImageJ (https://imagej.net/ij/).

### Coverslip IFA of late-stage parasites

Late-stage parasites were settled on to a poly-D-lysine coated #1.5 12 mm coverslip for 30 minutes. After removing the supernatant, the parasites were fixed with 4% paraformaldehyde for 10 minutes, then washed 3 times in PBS. Cells were permeabilized with 0.1% Triton X-100 in PBS for 10 min and washed three times in PBS for 3 min. Blocking solution (3% (w/v) BSA in PBS) was added and samples were incubated overnight at 4 °C. Primary antibodies were diluted in blocking solution and added to the coverslips for 2 hours at room temperature. Coverslips were washed 3 times for 3 minutes in PBS and then incubated with secondary antibody in blocking solution for 45 minutes. Coverslips were washed three times with PBS, then mounted onto a slide with VectaShield Vibrance with DAPI. Parasites were visualized on a Zeiss LSM980 with Airyscan2 for super-resolution microscopy. A 63x objective with a numerical aperture of 1.4 was used. Dilutions for primary antibodies were 1:200 for rat anti-HA and 1:300 for rabbit anti-*Pf*Actin.

### Ultrastructure Expansion Microcopy (U-ExM)

Late segmented schizont stage *Pf*Coronin^WT^-smHA or *Pf*Coronin^R100K/E107V^-smHA parasites were purified by Percoll gradient centrifugation. Various age groups of parasites, as specified in the results section, were utilized for U-ExM. For experiments aimed at confirming RBC content uptake, RBCs in a hypotonic solution were pre-loaded with biotin-dextran (ThermoFisher Cat. No. D1956), as described in the endocytosis assay above. The samples were treated with pre-warmed 4% paraformaldehyde-0.1% glutaraldehyde in PBS for 20 minutes at 37 °C and prepared for U-ExM following the procedure described previously^31,32^. The following antibodies were employed for visualization: rat anti-HA for *Pf*Coronin-smHA (1:25; Sigma Cat. No. 11867423001), Sytox Deep Red for DNA (1:1000; ThermoFisher Cat. No. S11380), rabbit anti-aldolase as a marker for parasite cytoplasm (1:2000; Abcam Cat. No. ab207494), Streptavidin-488 for parasites containing internalized biotin-dextran (1:250; Thermo Fisher Cat. No. S11223), and Alexa Fluor 405-conjugated NHS ester for protein density (1:200; ThermoFisher Cat. No. A30000). All secondary antibodies (anti-rat IgG Alexa fluor 488 Thermo Fisher Cat. No. A11006, anti-rabbit IgG Alexa fluor 555 Thermo Fisher Cat. No. A21428) were diluted 1:500 in PBS.

### Statistical analysis

All experiments were performed with at least three biological replicates, unless described otherwise. For volcano plots of proteins from *Pf*Coronin coIPs, *p*-values were computed using unpaired t-tests. Protein quantifications (immunoblot) were compared using paired t-tests. RSA assays were compared using unpaired *t*-tests with the Welch correction. Uptake of host cell contents (endocytosis assay) was compared using unpaired non-parametric Mann-Whitney tests. All statistics were calculated using GraphPad Prism. No statistical methods were used to predetermine sample sizes. Researchers were blinded for analysis of the endocytic assay (uptake of host cell contents) and for all RSA assays.

