## Supplemental files for "Cellular and molecular basis of *Pf*Coronin function in artemisinin resistance in *Plasmodium falciparum*"

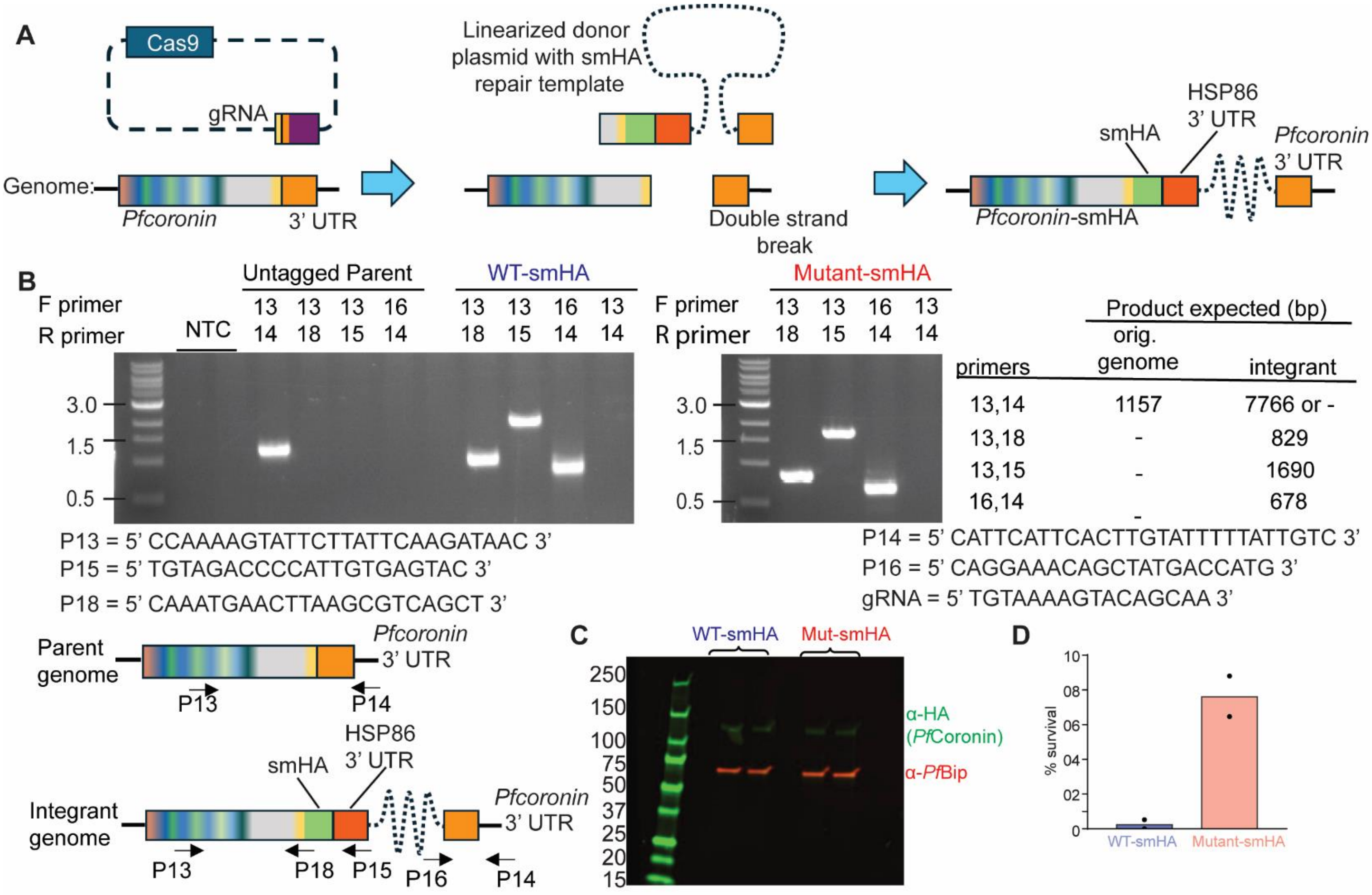


**Figure S1. Figure S1. Generation of spaghetti monster hemagglutinin epitope (smHA)-tagged WT and mutant transgenic lines.**

WT refers to parasites expressing wild-type *Pfcoronin*, and mutant refers to parasites expressing *Pfcoronin* containing the R100K/E107V mutations. Both lines were generated in the Pikine genetic background of *P. falciparum*.

A. Schematic for C-terminal endogenous tagging of WT and mutant parasites with smHA. Cas9 created a double-stranded break near the C-terminus of the *Pfcoronin* locus in either the WT or the mutant parent strain. This break was subsequently repaired via double crossover/homologous recombination between the *Pfcoronin* coding sequence (CDS; upstream of the break), the *Pfcoronin* 3' UTR, and matching sequences on the donor vector. This resulted in the integration of the entire donor vector, including an smHA tag appended to the C-terminus of *Pfcoronin.* B. PCR confirmation of smHA tagging in WT-smHA and mutant-smHA lines. Expected sizes from PCRs using genomic DNA templates from untagged (parental/ "orig. genome") *Pfcoronin* and *Pfcoronin-smHA* ("integrant") lines are detailed in the associated table. Primers used for the PCRs are shown and their binding positions are annotated in the schematic of the parasite genome. The gRNA sequence used is also shown. C. Confirmation of smHA-tagged WT and mutant lines via western blotting using anti-HA (mouse mAb 2-2.2.14; green) and anti-*Pf*BiP (loading control; rabbit polyclonal red) antibodies; samples were generated from late-stage parasites. Two biological replicates are shown for each strain. D. Tightly synchronized 3-hour-old ring-stage WT-smHA and mutant-smHA parasites were pulsed with 700 nM DHA for 6-hours, and survival was assessed 66 hours later. As expected, substantial survival (i.e., artemisinin resistance) was observed in the mutant-smHA line. Generally, the minimum survival threshold for ART resistance is 1%. Means from two independent biological replicates are shown.

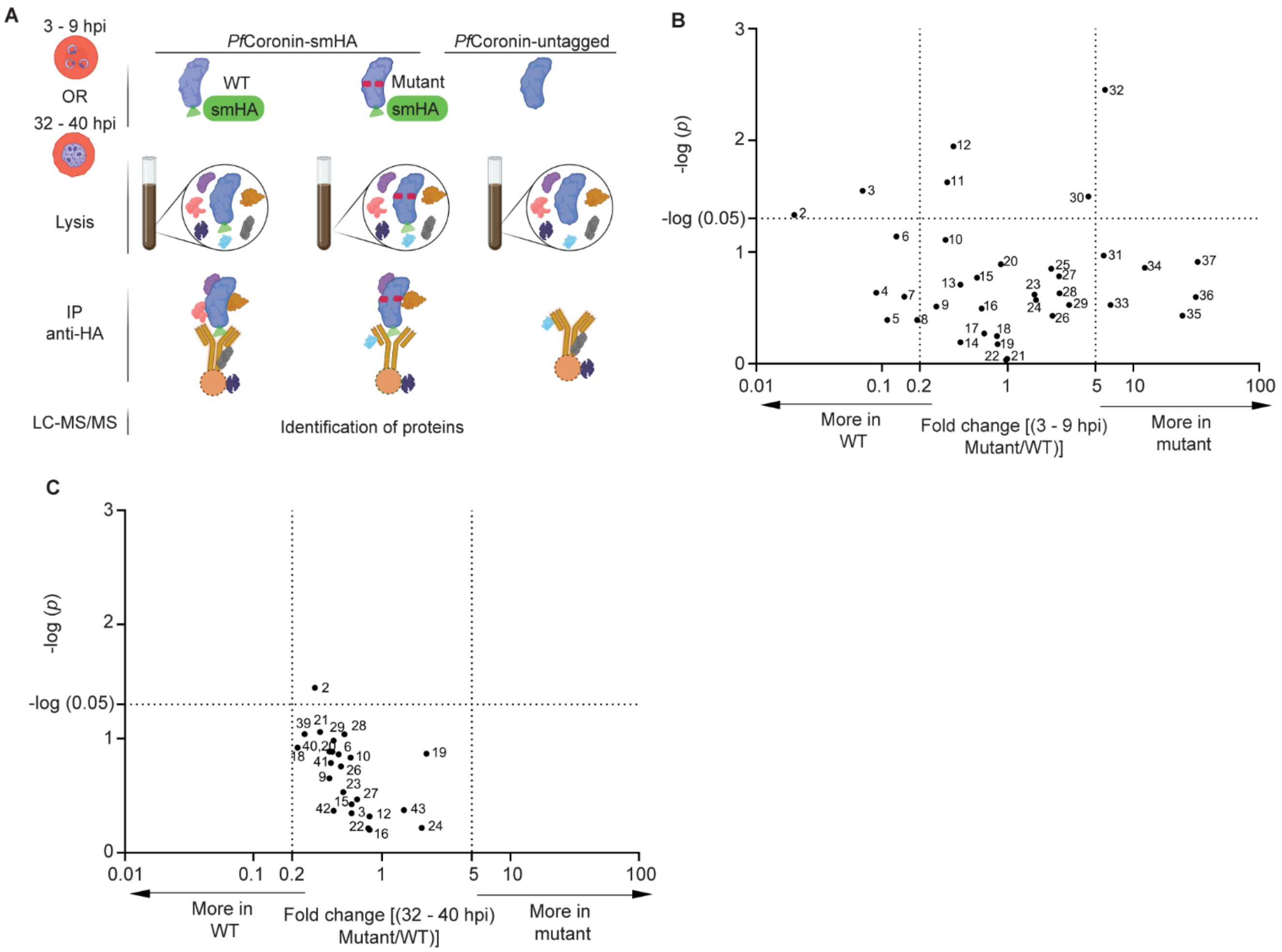


**Figure S2. Immunoprecipitation of *Pf*Coronin-smHA and associated proteins, related to Figure 1A-B and Tables S3-4.**

A. Immunoprecipitation of *Pf*Coronin-smHA and associated proteins from young ring-stage (3-9 h) and late-stage (32-40 h) parasites using anti-HA antibodies. Co-purified proteins were identified by LC-MS/MS, with an untagged parental control used to filter nonspecific interactions. hpi = hours post invasion. B-C. A volcano plot of hit interaction partners shows changes between *Pf*Coronin$^{WT}$-smHA and *Pf*Coronin$^{mutant}$-smHA in (b) rings, (C) trophozoites. The x-axis shows the fold change between *Pf*Coronin$^{WT}$-smHA and *Pf*Coronin$^{mutant}$-smHA bait proteins. The Y axis shows the statistical significance of the change between *Pf*Coronin$^{WT}$ and *Pf*Coronin$^{mutant}$ (unpaired t-test). The 37 hit proteins (for rings in B) and 24 hit proteins (for trophozoites in C) plotted on the volcano plot were found in all bioreps (either *Pf*Coronin$^{WT}$-smHA or *Pf*Coronin$^{mutant}$-smHA) and have at least 5% of the *Pf*Coronin protein abundance. In rings (B) two proteins (#1, *Pf*3D7_0517700, eukaryotic translation initiation factor 3 subunit B; and #38, *Pf*3D7_1424400, 60S ribosomal protein L7-3, putative) fulfilled these criteria but could not be displayed because they were detected only in *Pf*Coronin$^{WT}$-smHA (*Pf*3D7_0517700) or only in *Pf*Coronin$^{mutant}$-smHA pulldowns (*PF*3D7_1424400), so a log fold change could not be computed. A full list of hit proteins can be found in Table S3 (for rings) and Table S4 (for trophozoites). Proteins of interest are defined as either those with $p < 0.05$ (above the dotted line on the y-axis), or a 5-fold or greater change between *Pf*Coronin$^{WT}$-smHA and *Pf*Coronin$^{mutant}$-smHA parasites (to the left (0.2) or right (5) of the vertical dotted lines on the x-axis). The proteins are numbered and are detailed in Table S3 (for rings) and Table S4 (for trophozoites). Each data point represents two or three independent biological replicates for rings and trophozoites, respectively. WT and mutant indicate wild-type or R100K/E107V-mutant *Pfcoronin* in Pikine *P. falciparum* strain.

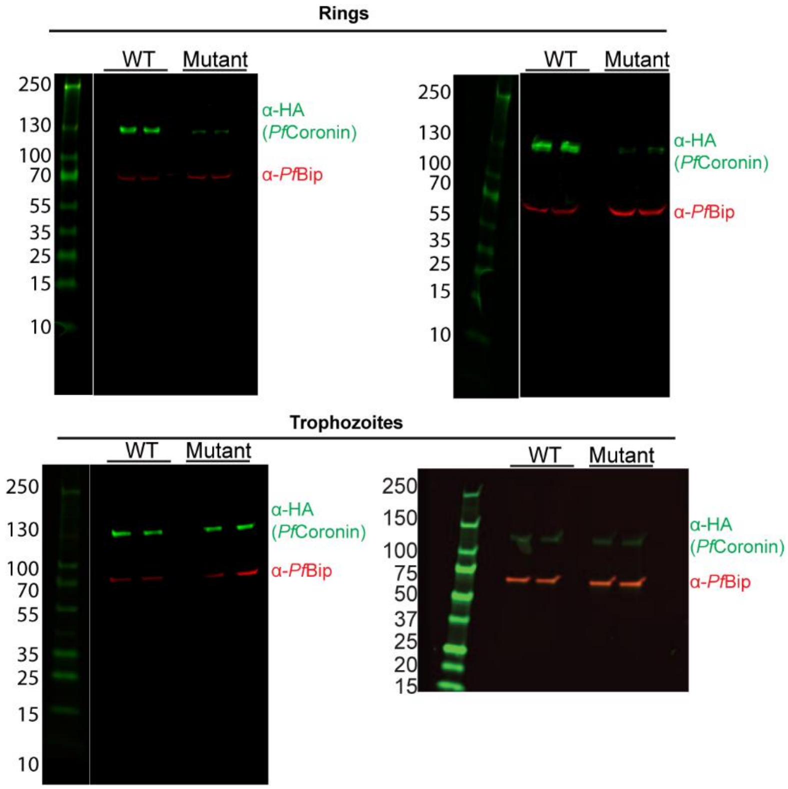


**Figure S3**. **Total *Pf*Coronin protein levels in parasite lysates, related to Figure 2A-B.**

Total *Pf*Coronin levels in lysates from parasites expressing *Pf*Coronin$^{mutant}$-smHA and *Pf*Coronin$^{WT}$-smHA were compared by western blot. Blots were probed with anti-HA (mouse monoclonal; green) and anti-*Pf*BiP (loading control; rabbit polyclonal, red) antibodies. All western blots corresponding to replicates quantified in Figure 2A and 2B are shown. WT = *Pf*Coronin wild-type smHA; mutant = *Pf*Coronin R100K/E107V smHA, both in Pikine *P. falciparum* strain.

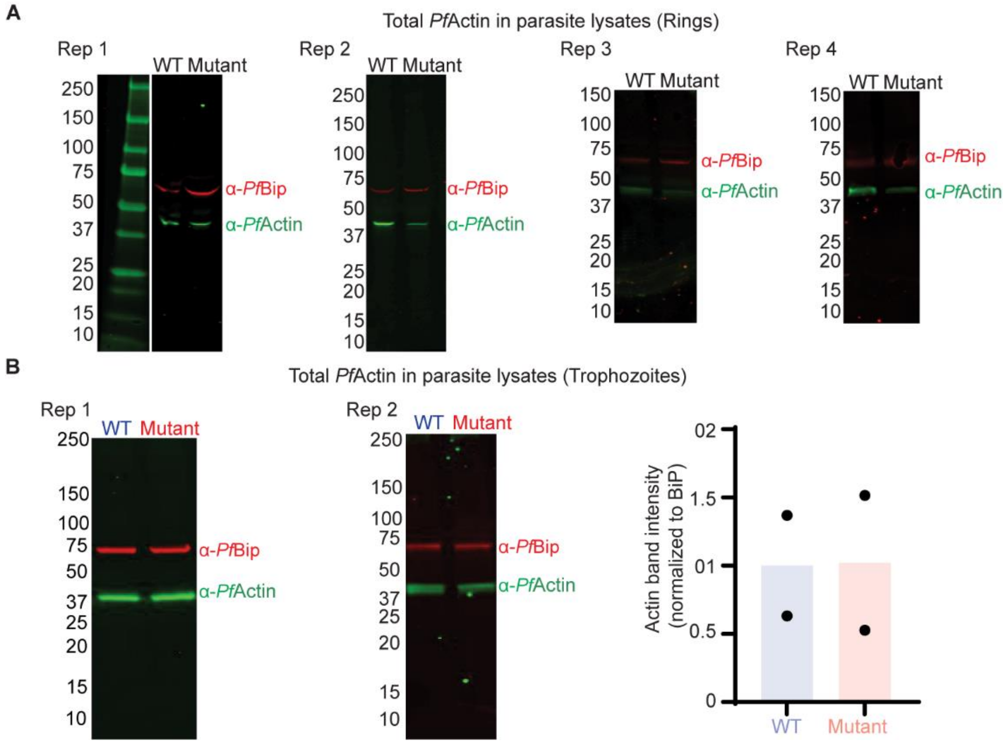


**Figure S4. *Pf*Actin protein levels in ring and trophozoite stages of PfCoronin$^{WT}$ and *Pf*Coronin$^{mutant}$ parasites (related to Figure 2D).**

WT and mutant indicate wild-type or R100K/E107V-mutant *Pfcoronin* expressed in the Pikine *P. falciparum* strain. smHA-tagged versions of *Pf*Coronin$^{WT}$ and *Pf*Coronin$^{mutant}$ were used.

(A) Western blot analysis of total *Pf*Actin in ring-stage parasites shows reduced *Pf*Actin levels in *Pf*Coronin$^{mutant}$ compared to *Pf*Coronin$^{WT}$. Blots were probed with anti-*Pf*Actin (mouse monoclonal; green) and anti-*Pf*BiP (loading control; rabbit polyclonal, red) antibodies. All western blots corresponding to replicates quantified in Figure 2D (n = 4) are shown.

(B) In late trophozoite-stage parasites, total *Pf*Actin levels are unchanged between *Pf*Coronin$^{mutant}$ and *Pf*Coronin$^{WT}$. Blots were probed as in (A). Quantification was based on two independent biological replicates.

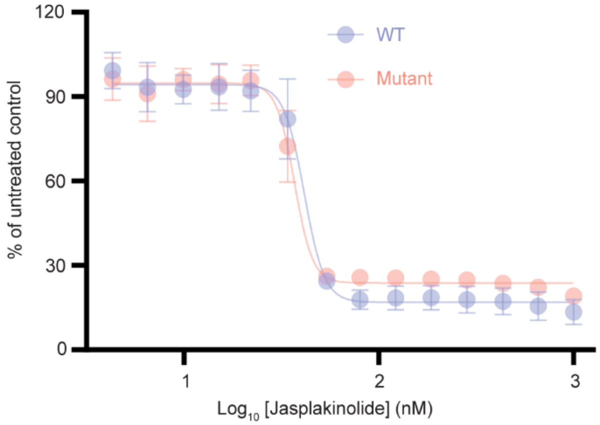


**Figure S5. Concentration-response assay using Jasplakinolide against both *Pfcoronin* WT and *Pfcoronin* mutant parasites, related to Figure 2E.**

A standard 72-h assay shows no difference between WT (Pikine parent; $EC_{50}$ = 40.1 ± 1.3 nM) and *Pf*Coronin mutant ((DHA selected Pikine_R line[1]; $EC_{50}$ = 37 ± 2.6 nM). Data points shown are average measurements from three independent biological replicates.

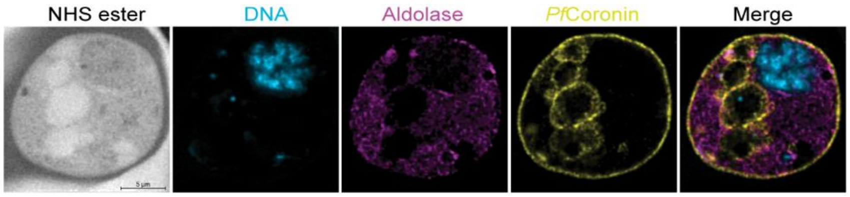


**Figure S6. Representative U-ExM image showing co-staining of *Pf*Coronin$^{WT}$-smHA and *Pf*Aldolase.**

A representative U-ExM image shows *Pf*Coronin$^{WT}$-smHA-labeled intracellular chains of small and large vacuoles in a trophozoite. Alexa Fluor 405-conjugated NHS ester indicates protein density, Sytox Deep Red marks DNA (cyan), Aldolase 555 stains parasite cytoplasm (magenta), and HA (*Pf*Coronin) appears yellow. Images are maximum-intensity projections.

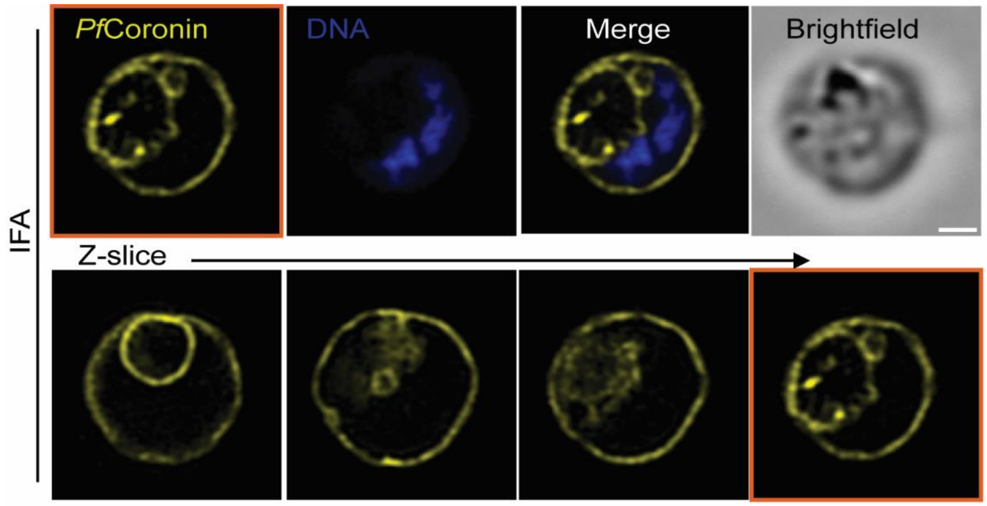


**Figure S7. Super-resolution fluorescence microscopy of *Pf*Coronin$^{WT}$-smHA in early schizonts.**

Super-resolution fluorescence microscopy (Zeiss LSM980 with Airyscan2) shows localization of *Pf*Coronin$^{WT}$-smHA in early schizont-stage parasites. Scale bar, 2 µm.

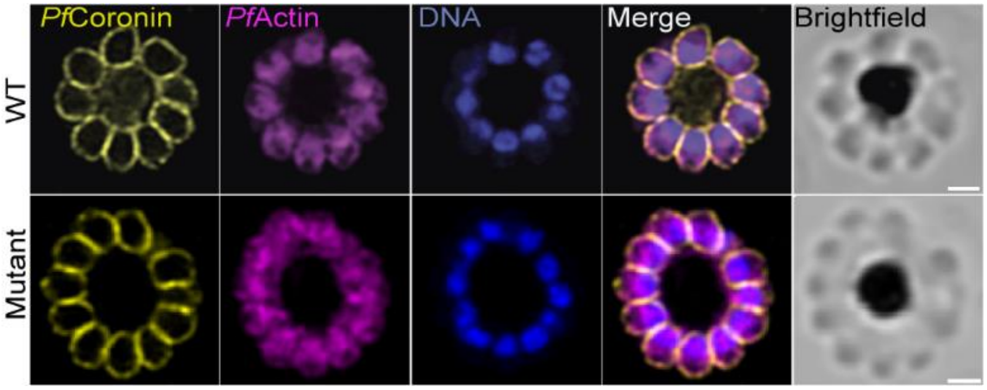


**Figure S8. Immunofluorescent characterization of *Pf*Coronin and *Pf*Actin localization**.

Super-resolution fluorescence microscopy (Zeiss LSM980 with Airyscan2) shows localization of *Pf*Coronin and *Pf*Actin in segmented schizonts. Scale bars = 2 µm. Parasites used were *Pf*Coronin$^{WT}$-smHA and *Pf*Coronin$^{mutant}$-smHA (mutant; R100K/E107V substitutions), both in the Pikine genetic background of *P. falciparum*.

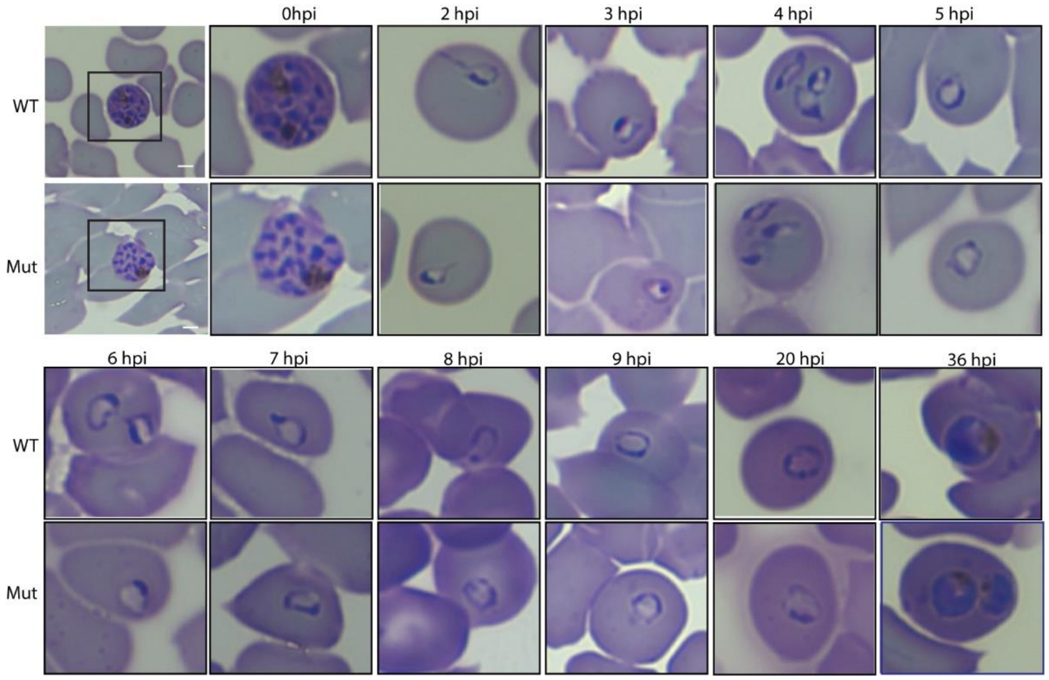


**Figure S9. Timecourse of Giemsa stained *Pf*coronin$^{WT}$-smHA and *Pf*Coronin$^{mutant}$-smHA parasites.**

Representative time-course images of Giemsa-stained, tightly synchronized (Percoll-purified) *Pf*Coronin$^{WT}$-smHA and *Pf*Coronin$^{mutant}$-smHA parasites tracking ring-stage development and transition to trophozoites. WT refers to *Pf*Coronin wild-type-smHA and mutant (mut) to *Pf*Coronin (R100K/E107V)-smHA, both in the Pikine *P. falciparum* background. Scale bar, 6 µm (first image); timepoints show zoomed views.

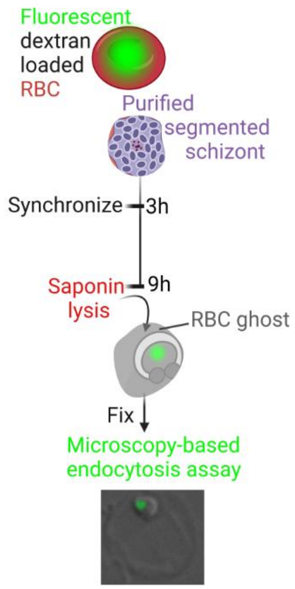


**Figure S10. Diagram of sample preparation for endocytosis.**

RBCs were lysed in a hypotonic solution containing fluorescent-dextran (Alexa 488). After re-sealing the RBCs (trapping the fluorescent dextran within), schizonts were added and allowed to re-invade for 3 hours. As the parasites (rings) grew within the fluorescent-dextran-containing RBCs, fluorescent -dextran was endocytosed and accumulated in the DV and other compartments. The parasites were then synchronized to remove any remaining schizonts and allowed to grow for a further 6 hrs, closely mimicking the RSA setup. RBC cell contents were removed by saponin lysis and parasites were fixed for microscopy. DIC and AF488 channel images were captured using a 100X oil immersion objective on a Zeiss Axio Observer. Representative image is shown. Images were blinded prior to analysis. Analysis was carried out for regions of interest (ROIs) where an RBC ghost could be visualized around a green channel signal in the merged image. Vesicle size (area), internalized fluorescence intensity (mean gray value), and integrated density (over the entire vesicle) of green-channel fluorescence signal was assessed.

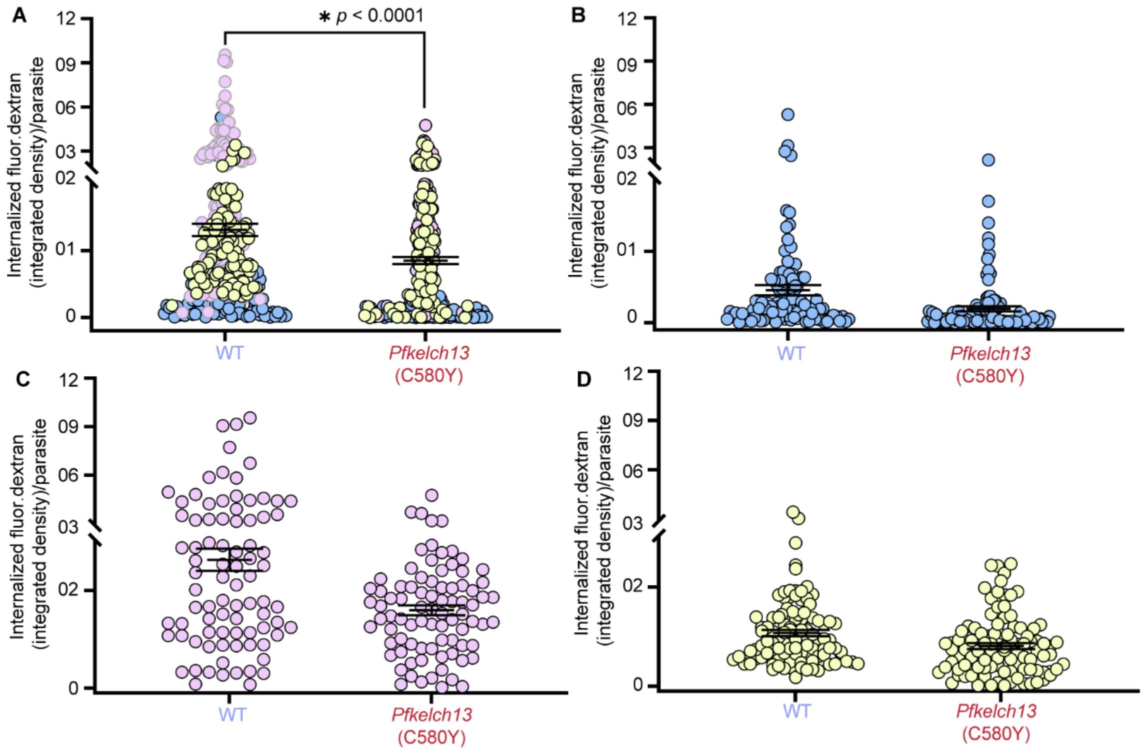


**Figure S11. Quantification of microscopy-based endocytosis assay comparing parasites carrying WT and mutant *Pfkelch13* (C580Y).**

Comparison of internalized fluorescent-dextran in WT (Pikine parent) vs. *Pfkelch13* (C580Y, Pikine parent) parasites by (A) integrated density (vesicle size x intensity). WT, n = 286; *Pfkelch13* (C580Y) n = 286. Each point represents uptake in a single parasite. *p*-values (Mann-Whitney test) and percent reduction of the mean are indicated. Data are pooled from the biological replicates shown in (B)–(D).

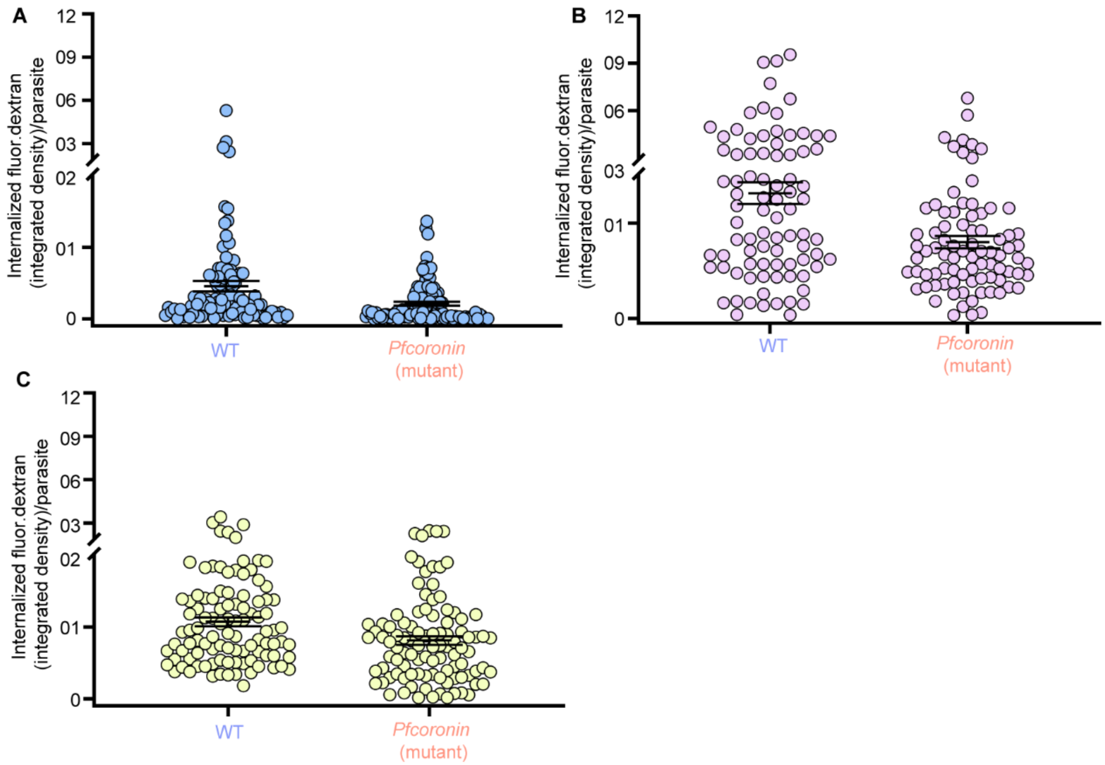


**Figure S12. Quantification of microscopy-based endocytosis assay comparing parasites carrying WT and mutant *Pfcoronin* (R100K/E107V), related to Figure 5.**

(A–C) Internalization of fluorescent dextran in *Pfcoronin* WT (Pikine parent) and *Pfcoronin* mutant (R100K/E107V, CRISPR-edited Pikine line) ring-stage parasites (3–9 hours post-invasion) is shown for individual biological replicates, which were pooled for analysis in Figure 5. Each point represents uptake by a single parasite.

**Table S5. List and characteristics of *Plasmodium falciparum* cell lines used in this study.**

| Purpose in Study | Parasite Line | Genetic Background | Details | SNP(s) in *Pf*coronin or *Pfkelch13* | smHA-tagged |
|---|---|---|---|---|---|
| **RSA timecourse, JAS survival experiments** | Pikine parent (SenP019.04); also called *Pfcoronin*$^{WT}$ | Pikine | Culture-adapted *P. falciparum* line from Senegal[8] | No | No |
| | Pikine_R | Pikine | DHA selected resistant line[8] | *Pfcoronin* R100K/E107V | No |
| **Endocytosis experiments** | Pikine parent (SenP019.04); also called *Pfcoronin*$^{WT}$ | Pikine | Culture-adopted *P. falciparum* line from Senegal | No | No |
| | *Pfcoronin*$^{R100K/E107V}$ | Pikine | CRISPR-edited line (cF5) | *Pfcoronin* (R100K/E107V) | No |
| | *Pfkelch13*$^{C580Y}$ | Pikine | CRISPR-edited line | *Pfkelch13* (C580Y) | No |
| **All other experiments (microscopy, pulldowns)** | *Pfcoronin*$^{WT}$*-smHA* | Pikine | CRISPR-edited line | No | Yes |
| | *Pfcoronin*$^{R100K/E107V}$*-smHA* | Pikine | CRISPR-edited line (cF5) | *Pfcoronin* (R100K/E107V) | Yes |
